# Bidirectional network hubs: NT-genes as optimal targets for partial cancer reversal

**DOI:** 10.64898/2026.04.27.721122

**Authors:** Gabriel Gil, Rolando Perez, Augusto Gonzalez

## Abstract

**Background:** The complexity of gene regulatory networks, involving thousands of genes, poses a fundamental challenge to understanding cancer phenotype reversal. However, recent evidence suggests that the effective dimensionality of normal and tumor transcriptional manifolds is low, and that small panels of genes can discriminate perfectly between normal and tumor samples.

**Methods:** We build upon two previously developed concepts: (i) highly accurate normal and tumor gene markers (namely, N-and T-markers), defined as genes with exclusive expression intervals in normal and tumor samples, respectively; and (ii) gene deregulation networks (GDNs), represented as directed acyclic graphs encoding causal relationships between gene deregulation events. A subset of genes appearing in both marker classes (NT-markers) act as bridging nodes between the N-and T-GDNs. Starting from these elements, we introduce a quantitative dynamical model based on node frequency and connectivity to assess how gene intervention effects propagate through the GDN and thereby predict their overall impact on the tumor tissue.

**Results:** According to the model, interventions on *pure* T-markers (T-markers that are not NT-markers) produce effects largely confined to the T-GDN, with a minimal perturbation of the N-network. Interventions on *pure* N-markers (N-markers that are not NT-markers) generate a perturbation of both networks, but with limited effect. In contrast, interventions on NT-markers with high activation frequency in both tumors and normal state (e.g., *AGER* in lung adenocarcinoma: 98% in tumor samples, 75% in normal samples) can induce bidirectional phenotype shifts. For an effective combination of targets, coverage across tumor samples must be maximized. At the same time, in the T-GDN the number of nodes unreached by the reverse cascade following the intervention must be minimized, as these regions may act as escape routes for the tumor. Escape probability further depends on the tumor stage and the tumor’s activation rate of new T-genes. When targeting NT genes, high frequency in normal samples should also be prioritized.

**Conclusions:** High-frequency NT-genes, due to dual network connectivity and tissue relevance, represent optimal targets for achieving at least partial phenotype reversal. This framework provides a quantitative guide for prioritizing gene therapy targets and designing combination strategies that balance coverage, escape minimization, and normal tissue relevance.

## 1. Introduction

The reversion of the tumor phenotype to a normal cell program is a dream that originated decades ago [1-3] and has recently gained a second wind [4-6]. The question of whether manipulating a small set of genes can reverse the tumor phenotype appears counterintuitive given that gene regulatory networks [7,8] involve thousands of interacting nodes. However, the highly interconnected architecture of these networks implies that the expression of many genes varies in a coordinated manner rather than independently, so that modulating the expression of a limited number of genes may suffice to induce widespread transcriptional—and consequently phenotypic— changes. Evidence of this underlying co-variation is provided by the surprisingly low effective dimensionality of the normal and tumor transcriptional states. Indeed, geometric analyses of gene expression manifolds indicate that the intrinsic dimension of these biological states is typically less than 10 [9,10]. This reduced dimensionality suggests that the essential differences between normal and cancerous states might be captured by a relatively small number of key regulatory genes, which are natural candidates for phenotype reversal interventions.

Identifying target genes for cancer therapy has long been a central goal in omics research, particularly through the analysis of large-scale RNA-Seq data from normal and tumor samples. Recent efforts in this direction have revealed the existence of genes that serve as markers of the normal and tumor states (namely, N-and T-markers or N-and T-genes) [11]. Importantly, the concepts of N-and T-markers generalize those of tumor suppressors and oncogenes [12] to capture broader patterns of cancer-related gene expression deregulation [11,13]. Remarkably, small panels of such genes—often fewer than ten—can perfectly discriminate between normal and tumor samples across multiple cancer types [11]. These findings point to the possibility that genes in such panels could play a crucial role in phenotype reversal.

Parallel work on deregulation networks has introduced a conceptual framework distinguishing the regulatory architectures of normal tissues and tumors [14]. By construction, normal tissues predominantly activate N-genes within an N-network, while tumors activate T-genes within a T-network. Somatic evolution in normal tissue involves the progressive deregulation of the N-network, whereas clonal evolution in tumors operates within the T-network [13,14]. Within this framework, phenotype reversal—the transition from a tumor back toward a normal state— necessarily involves both networks simultaneously.

Genes that appear as markers in both normal and tumor samples, termed NT-genes or NT-markers, emerge as particularly promising candidates for therapeutic intervention [15]. Because NT-genes participate in both networks, they may influence both the suppression of T-gene activity and the reactivation of N-gene programs. Although not explicitly conceptualized as NT-markers, experimental studies across multiple tissues suggest that top-ranking genes from this class can induce partial phenotype reversal both in vitro and in vivo [16-25].

The present work aims to develop a minimal quantitative model that predicts the outcomes of genetic interventions based on the N-and T-genes (specially the NT-genes) and the corresponding deregulation networks. By incorporating simple quantitative measures—specifically, the activation frequency of markers in normal and tumor samples and the fraction of the T-network reachable by intervention-induced deactivation cascades—we seek to provide a practical guide for prioritizing gene therapy targets. Interventions on specific genes can be readily simulated using previously obtained deregulation networks [14]. The model generates distinct predictions for interventions targeting each gene class—predictions that can be tested against new or existing experimental data. Ultimately, we aim for this framework to contribute to rational design of gene therapies targeting partial and complete reversal of the cancer phenotype.

## 2. Methods

### 2.1. Definition of N-, T-, and NT-genes

Following the procedures established in [11] and [13], we begin with bulk RNA-Seq data from The Cancer Genome Atlas (TCGA) [26-28], including gene expression profiles of normal and tumor samples for each cancer type. A gene is classified as an N-marker if there is a normal-exclusive expression interval populated by a significant fraction of normal samples. Conversely, a T-marker is characterized by the presence of a significantly populated tumor-exclusive expression interval. Statistical significance is established by setting a minimum percentage of samples in the exclusive interval (e.g., 10% for tumor samples). A gene qualifies as an NT-marker if it belongs simultaneously to both the N-and T-marker classes. NT-genes are of particular interest because they represent nodes that are functionally relevant in both physiological and pathological states and may therefore serve as bridges between the N-and T-deregulation networks [13,14].

We map the continuous expression levels of N-, T-and NT-genes to the following states: “N-active” (−1) if the expression lies within the normal-exclusive interval, “T-active” (1) if it lies within the tumor-exclusive interval, and “inactive” (0) otherwise. We refer to an N-gene as *pure* if it can assume only the N-active and inactive states. Likewise, a T-gene is termed *pure* if it can assume only the T-active and inactive states. This contrasts with NT-genes, which can be N-or T-active, although not necessarily inactive.

### 2.2. Marker Activation Frequency

A fundamental property of any marker is its activation frequency within the corresponding sample set. Typically differing N-and T-activation frequencies are both central to assessments involving NT-genes. For example, in prostate adenocarcinoma (PRAD), the NT-gene *SH3GLB1* shows frequencies f_N_ = 0.25 and f_T_ = 0.58, indicating that a possible tumor test based on that gene is sensitive but not quite specific.

We interpret a high activation frequency as indicating relevance for tissue identity [14]. A gene activated in 98% of tumors, such as *AGER* in lung adenocarcinoma (LUAD) is likely to be performing essential functions within the tumor state; conversely, a gene activated in 81% of normal samples, such as *STX11* in LUAD, is likely to be important for normal tissue physiology. This interpretation has direct implications for therapeutic intervention: targeting a gene with high f_T_ should produce substantial damage to the tumor, while targeting a gene with high f_N_ might disrupt homeostasis.

T-genes with low activation frequencies, on the other hand, exhibit high deregulation capacity [14], that is they may initiate extensive deregulation cascades. For N-genes, the deregulation process corresponds to their deactivation. Thus, genes with high f_N_ are also the initiators of deregulation cascades.

Conversely, intervention on high frequency T-genes or low-frequency N-genes, may lead to extensive reverse cascades, propagating through their respective networks according to rules specified in [14].

### 2.3. Gene Deregulation Networks

Gene deregulation networks (GDNs) were inferred in [14] from expression data using probabilistic causal discovery algorithms. The N-and T-GDNs describe the dynamics within the normal tissue and tumor, respectively. The transition from normal tissue to tumor involves both networks. These GDNs provide the substrate on which spontaneous evolution unfolds and perturbations—such as the forced activation or deactivation of nodes—propagate. A set of rules was established in [14] in order to make this dynamics explicit in each case.

Here, we focus on the effects of a gene interventions in a tumor tissue. The rules from Ref. [14] applying in this context are the following:

- **r8**. Forced activation of an N-gene in a tumor triggers an activation cascade of N-genes with higher activation frequencies.
- **r9**. Forced deactivation of a T-gene in a tumor triggers a deactivation cascade of T-genes with lower activation frequencies.
- **r10**. Forced N-activation of an NT-gene in a tumor triggers an activation cascade of N-genes with higher N-frequencies and a deactivation cascade of T-genes with lower T-frequencies.

A simple Monte Carlo algorithm was designed to simulate the evolution dynamics. In the case of intervention of pure T-genes in a tumor initial state, for example, the algorithm proceeds as follows. The system is initialized with a subset of T-genes active. This initial state may be taken from a tumor sample in the dataset, for example.

The intervention is then applied to the set of genes. It means that two processes will be running in the network. Deactivations of T-genes start from the intervened genes and propagate along the transposed T-GDN, whereas new activations of T-genes start from the already active genes and propagate along the edges of the T-GDN. In principle, the two processes act at different rates.

Genes in the reverse cascade following intervention and active genes in the tumor are randomly sampled per iteration. An additional assumption is taken that once a gene is deactivated it can not be activated again.

Similar evolution dynamics apply in other cases. Rules r1-r10 of [14] and the corresponding GDNs are used when they are needed.

## 3. Predictions, Results and Discussion

### 3.1 Qualitative picture of single-gene interventions

Figure 1 qualitatively represents the fate of a tumor following intervention on a single gene. Panel (a)depicts the initial tumor state: only T-markers (including NT-genes) are activated, represented as circles with upward arrows; deactivated genes (N-markers) lack arrows. In the following three panels, the intervened gene is marked in blue, and causal relations are indicated by red lines. The diagram emphasizes that T-genes typically outnumber N-genes, and that NT-genes constitute the interface between the two networks.

**Figure 1:**
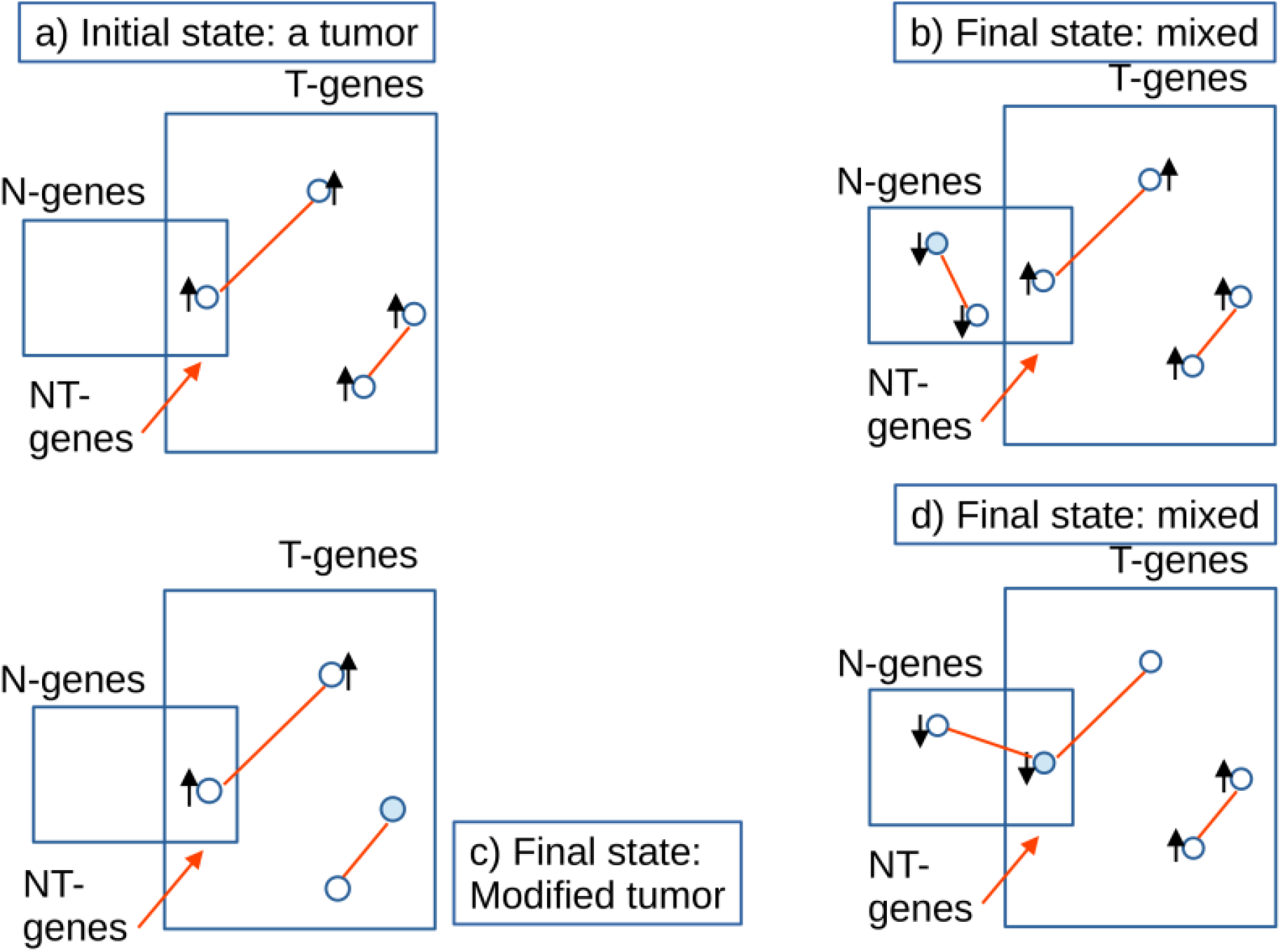
Schematic representation of the possible outcomes of a single-gene intervention in a tumor tissue. **(a)** The initial state. **(b)** Intervention of an N-gene. **(c)** Intervention on a T-gene. **(d)** Intervention on an NT-gene. The intervened gene is marked in blue. The activation state of genes is represented with arrows, whereas causal relationships are indicated by red lines.

Three distinct outcomes are possible depending on the class and network position of the intervened gene:

- **Case (b):** An intervention that activates a pure N-gene produces a mixed state in which some N-markers are newly activated, and some T-markers remain active. We define this outcome as partial phenotype reversal of the tumor—the cell population now exhibits features of both normal and tumor states. Rule **r8** prescribes a cascade in the N-network; however, deactivations of T-genes could be indirectly induced through NT-genes (not shown). Even if T-genes are not affected by the intervention, a mixed state still arises due to the coexistence of induced N-activations, and pre-existing T-activations.
- **Case (c):** An intervention that deactivates a pure T-gene, according to rule **r9**, affects only the active T-markers. The result is a modified tumor state, not a reversal. This case likely corresponds to the typical outcome of many targeted therapies, which suppress oncogenic signaling without restoring normal differentiation programs. In principle, there could be some N-markers indirectly activated through NT-genes (not shown). Rule **r9** does not explicitly account for this possibility. Unlike the case (b), where a mixed state appears unconditionally, in this scenario a mixed state occurs only if the pathway toward the N-GDN through NT-genes is effective.
- **Case (d):** The intervention on an NT-gene produces a mixed state in which some N-markers are newly activated and some T-markers are deactivated. According to rule **r10**, the NT-genes directly induce cascades in both networks. Unlike the case (b), the mixed state in this case is achieved by counteracting the spontaneous deregulatory dynamics of both the normal and tumor states.

Thus, the central prediction of this qualitative picture is that NT-genes are uniquely positioned within both networks to induce phenotype reversal. The effect of an intervention on pure N-and T-genes, on the other hand, is expected to be more confined to their respective networks and therefore less prone to trigger robust phenotype reversal.

### 3.2 Multiple T-targets

In this section, we consider combining interventions on multiple T-genes. The motivation for using multiple targets is twofold: on the one hand, to increase coverage across tumor cases, and on the other, to reduce the escape routes of the tumor, that is, network pathways through which it can persist despite intervention.

The fraction of tumor samples covered by the intervention on a set of T-targets is defined in terms of the composed frequency:

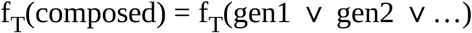

that is, the fraction of tumor samples in which at least one target gene is active (the v symbol is the logical OR). This quantity coincides with the sensitivity (true positive rate) of a tumor test based on a gene panel comprising gen1, gen2, etc., and provides a conservative estimate of the fraction of tumors potentially affected by the combined intervention. A value f_T_(composed) = 1, corresponding to an optimally discriminating panel [11], means that all possible tumor states are directly perturbed by the intervention.

### 3.3 Escape under single-gene interventions

Tumors are dynamical systems that evolve over time. Even after a successful intervention, residual tumor cells may harbor pathways that enable escape and disease progression. We accounted for this process in the simulations of Ref. [14] by letting the tumor evolve at a given rate according to its spontaneous dynamics. This dynamic, together with the fact that not all T-genes are reached by propagating the effects of an intervention, give rise to what we refer in that paper to as escape routes of the tumor. Indeed, unreached genes become seeds for new spontaneous deregulation cascades, ultimately leading to the re-emergence of the tumor phenotype.

Rather than performing a simulation of the dynamics, we may inspect the structural features of the network to illustrate how they can foster escape routes for the tumor. In particular, in Figure 2, we evaluate the fraction of the network that remains unreached by a deactivation cascade starting at the highest-frequency T-gene. The x-axis is the depth of the network from the cascade initiator measured in steps. At the first step, we only reach the parents of the intervened gene; at the second step, only the parents of those parents, and so on.

**Figure 2:**
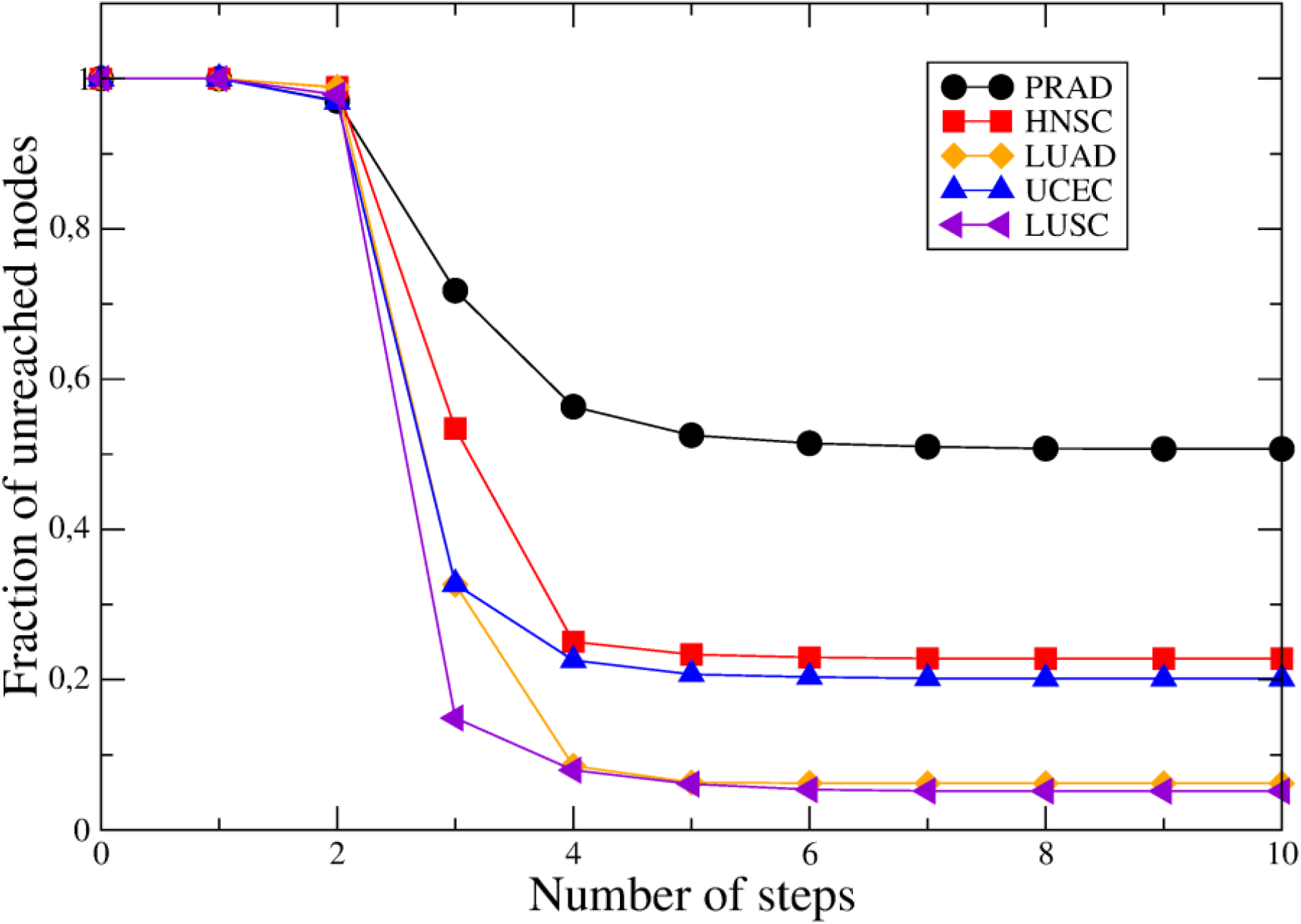
Fraction of nodes in the T-network unreached by the reverse cascade following the intervention on the highest-frequency T-gene. Five tumors are considered: PRAD, head and neck scamous carcinoma (HNSC), LUAD, uterine corpus endometrial carcinoma (UCEC) and lung scamous carcinoma (LUSC).

The data plotted in Figure 2 are based on the T-GDNs associated with the five cancer types studied in [14], that is prostate adenocarcinoma (PRAD), head and neck squamous carcinoma (HNSC), lung adenocarcinoma (LUAD), uterine corpus endometrial carcinoma (UCEC), and lung squamous cell carcinoma (LUSC). For prostate cancer, the most heterogeneous cancer, the cascade reaches almost 50% of the network, but for the two cases of lung cancer considered—adenocarcinoma and squamous cell carcinoma—the cascade reaches about 90% of the network. Notice that these fractions correspond to the maximally wide and maximally deep cascade, and that in the calculation all nodes are treated on equal footing despite of the fact that, as mentioned above, we understand that high-frequency nodes are more relevant for tissue identity.

Similar results may be obtained for a combination of targets. The simultaneous optimization of the combination frequency and of the fraction of the network reached by the intervention should be a central goal of therapeutic design.

### 3.4 Multi-target interventions in extreme cases: PRAD and LUAD

Prostate adenocarcinoma is one of the most heterogeneous tumors. This property is probably related to the fact that the normal and tumors attractors are relatively close [10,29]. As a consequence, the number of genes in the perfect panel is relatively large, 8 genes [11]. In Ref. [14], we performed simulations for the intervention on all of the panel genes, which nevertheless resulted in tumor escape. Here, we appeal to the procedure described in the previous section to estimate the escape probability of such an intervention in terms of the fraction of nodes unreached by the common maximal cascade initiated at the panel genes. The result is around 30%, as shown in Figure 3, top panel. Thus, the inclusion of 7 genes in addition to *EPHA10* allows one to cover the whole population of tumors, while reaching only 70% of the T-GDN.

**Figure 3:**
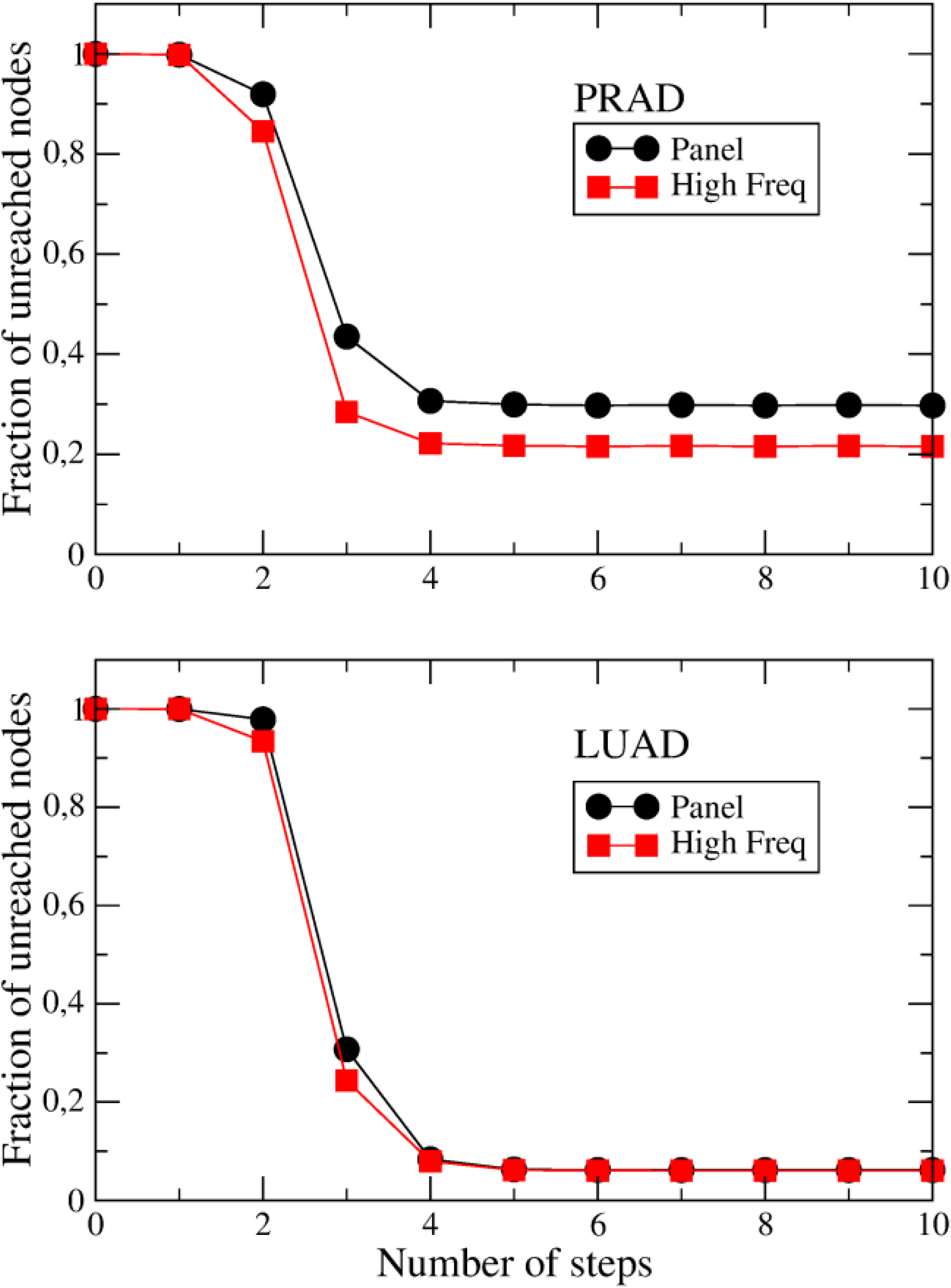
Fraction of nodes in the T-network unreached by the reverse cascade following the intervention on two gene sets. **(a)** The perfect panel of 8 genes and the set of 8 genes with the highest frequencies in PRAD. **(b)** The perfect panel of 2 genes and the 2 genes with the highest frequencies in LUAD.

In the same figure, we show results for interventions on the 8 genes with the highest frequencies in the tumor population. The composed frequency for this set of genes is only 0.69, but it reaches almost 80% of the network, leaving 20% of nodes as potential escape routes. Hence, by sacrificing full coverage, we are able to reduce the escape probability. This indicates that a compromise between coverage and therapeutic effectiveness—estimated in terms of transposed T-GDN traversability— will be required in the prostate cancer.

The bottom panel of Figure 3 shows similar results for lung adenocarcinoma (LUAD). The larger distance between attractors in this cancer type is linked to lower tumor heterogeneity, the presence of high-f_T_ genes and smaller perfect panels [11,14]. In this case, 2 genes are enough to discriminate between the normal tissue and the tumor: *AGER* and *SLC37A4*. The corresponding frequencies are 0.98 and 0.84. Substituting *SLC37A4* by the second-highest gene in the f_T_ ranking has little impact on the coverage (> 98%) or on escape probability (around 6%). Therefore, we prefer to consider the panel genes, which are related to independent pathways for tumor progression [14], as candidate targets for therapeutic interventions.

A fourth characteristic of tumors with distant attractors is that NT-genes are among the first in the high f_T_ and high f_N_ rankings (*AGER* in LUAD, for example). We will explicitly consider NT-genes in section 3.6.

### 3.5 Escape probability, tumor stage and T-activation rate

The fraction of the T-network that remains unreached by the reverse cascade sets a lower bound on the set of genes that the tumor may use as escape routes. The purpose of this section is to explicitly outline two additional factors that determine escape after intervention.

Let us recall the simulations presented in [14] for interventions in PRAD samples. In that study, the observation of a set of nodes with rising average activation was qualitatively interpreted as a sign of escape. Here we emphasize that the number of active T-genes — defined as genes with a mean activation greater than 0.5 — offers a quantitative perspective on escape. This number was linked in Ref. [13] to the degree of clonal evolution (malignancy) of the tumor. We ran new simulations according to the procedure described in Section 2.3. The deactivation rate was set to 0.5.

First, we note that the variable to be compared with the fraction of unreached nodes is the number of active nodes in the initial tumor sample, which defines the initial tumor stage. Consider, for example, an intervention targeting all genes in the perfect panel in PRAD. The nodes unreached by the reverse cascade represent only 30% of the network, i.e., about 1,800 nodes. These are the nodes that remain accessible on very long time scales. If the initial state is an evolved tumor, such as the one used in Ref. [14] with 2,640 active T-genes, the mean number of active genes can only decrease over time. Thus, evolved tumors can only become less malignant after an intervention that reaches a significant fraction of the T-network. Younger tumors are more plastic, in the sense that they have more possible ways to evolve.

The third factor affecting escape is the competition between deactivation and activation processes. This competition is more apparent in a younger tumor. To highlight this factor, we blocked the entire panel of genes and chose as a starting point a sample with 403 active nodes. We kept the deactivation rate at 0.5 and performed simulations for two values of the activation rate: 0.1 and 0.5. The results are shown in Figure 4.

**Figure 4:**
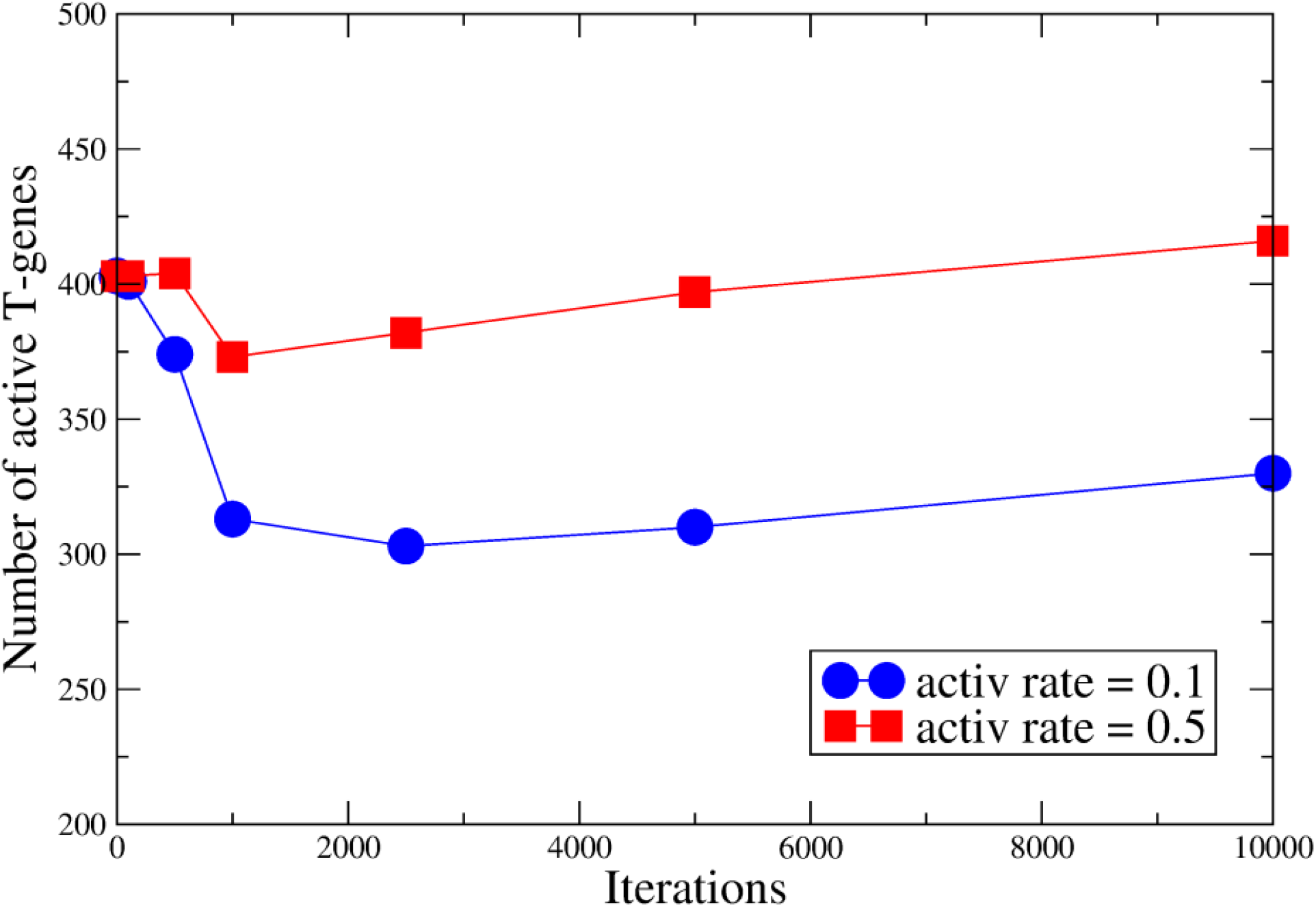
Pseudotime evolution of the number of active nodes following intervention on a sample with 403 active T-markers. The whole perfect panel of 8 genes is forcibly deactivated. The deactivation ratio in the reverse cascade is set to 0.5, whereas two values for the activation ratio of new T-markers are considered: 0.1 and 0.5.

Let us define the recovery time as the number of iterations at which the number of active genes equals the initial number of active genes. In the example, when the activation rate is high, the tumor recovers after approximately 10,000 iterations. For low activation rates, however, the recovery time is much longer. Thus, controlling the activation rate provides a means of achieving biological control over the tumor.

In conclusion, three quantities define tumor escape after intervention: the fraction of the network reached by the intervention, the tumor stage, and the activation rate. Controlling the last of these quantities should also be a central goal of the intervention.

### 3.6. NT-genes and tumor phenotype reversal

Intervention on the receptor for advanced glycation end-products (*AGER*) in lung adenocarcinoma (LUAD) cell lines provides a validation of the prediction of the qualitative picture, Figure 1d). This normal-above tumor-below gene exhibits near-perfect NT characteristics: f_T_ = 0.98 (activated in 98% of tumor samples) and f_N_ = 0.75 (activated in 75% of normal samples). Its high frequency in both states indicates that *AGER* is relevant for tissue identity in both normal lung and lung tumors.

Wang et al. [7] experimentally investigated the role of *AGER* in non-small cell lung cancer H1299 cells. Using lentiviral overexpression vectors, they increased *AGER* expression and observed profound phenotypic consequences: cell proliferation was significantly inhibited, migration and invasion capacities were reduced, and apoptosis was induced. Critically, these changes were accompanied by alterations in the expression of genes associated with normal lung epithelial differentiation. The authors concluded that *AGER* functions as a tumor suppressor in this context and its overexpression partially restores a more normal-like phenotype. Notably, Wang et al. also performed the reverse experiment, knocking down *AGER* in normal lung epithelial cells, which induced tumor-associated phenotypic changes. This bidirectional effect confirms that *AGER* functions as a master regulator capable of shifting cells between normal and tumor-associated states, exactly the behavior predicted for high-frequency NT-genes.

The effect of deactivating *AGER* on the T-network was already computed in our framework, as shown in Figure 2. The reverse cascade for LUAD starts precisely from *AGER*, which is the highest frequency T-gene. 90% of the T-network is reached by the reverse cascade.

With regard to the N-network, we first note that from the 492 N-genes in LUAD, 387 belong to the NT-category and, thus, they are indirectly subject to a change of activation state as a result of the reverse T-cascade. The reverse cascade induced in the N-network, according to rule **r8**, advances in the direction of higher f_N_, that is only *STX11* may be activated because it is the only gene with f_N_ higher than *AGER*’s frequency.

In conclusion, the intervention may reach 98% of tumor samples. The reverse cascade reaches 90% of the T-network (10% of escape possibilities) and may indirectly change the activation state of 70% of N-genes (those that belong to the NT category). The reverse cascade in the N-network comprises only two genes, but they are the two most relevant genes for the identity of the normal tissue.

We may analyze also the reverse experiment, i.e. knocking down *AGER* in a normal tissue. Rules **r3** and **r6** apply to the present context [14]. In this case there is no direct cascade in the T-network because *AGER* is the highest f_T_ gene. It is a single-gene cascade, but contains the most relevant gene for the tumor. The deactivation cascade in the N-network, on the other hand, acts in the same direction of the spontaneous process, but most probably at an accelerated rate. We may compute the maximal deactivation cascade in the same way as we computed the cascade in the T-network. The result is that the direct cascade reaches also 90% of the N-network. This is a huge number because commonly healthy normal tissue shows 60-70% of activated genes [13]. Indirectly, more than 300 NT genes may translate their change of activation state to the T-network.

In general, multi-target therapy with NT-genes should optimize, besides the fraction of tumors reached by the intervention and the escape probability, the relevance of targets in the N sector. A method for selecting the optimal targets was presented in [15] based on multi-objective (Pareto) optimization [30].

## 4. Concluding Remarks

The quantitative model presented here offers a simple yet powerful framework for predicting the outcomes of genetic interventions in cancer. By classifying genes into pure N-markers, pure T-markers, and NT-markers based on their expression patterns across normal and tumor samples, and by incorporating activation frequencies, we can anticipate whether an intervention will produce network-restricted effects or trigger a broader phenotypic shift involving both normal and tumor-associated programs. Activation frequency is not merely descriptive; it carries functional meaning. High-frequency genes are likely central to tissue identity in their respective states.

The central insight from our analysis is that NT genes occupy a privileged position at the interface of normal and tumor deregulation networks. Because they are functionally relevant in both physiological and pathological states, interventions targeting NT genes can simultaneously suppress tumor-specific programs and reactivate normal differentiation programs — the dual requirement for true phenotype reversal. This prediction, qualitatively illustrated in Figure 1d, is strongly supported by the *AGER* example in lung cancer [16]. With f_T_ = 0.98 and f_N_= 0.75 in LUAD, *AGER* is a near-perfect NT marker. Experimental overexpression of *AGER* in H1299 non-small cell lung cancer cells inhibited proliferation, migration, and invasion, induced apoptosis, and partially restored a normal-like epithelial phenotype. Critically, knockdown of *AGER* in normal lung epithelial cells induced tumor-associated changes, demonstrating bidirectional phenotypic control.

For combination therapies, the framework provides explicit quantitative guidance. The joint frequency: f_T_(gen1 v gen2 v…) estimates the fraction of the tumor population potentially affected. Simultaneously, the fraction of the T-network unreached by the reverse cascade (Figures 2 and 3) sets an upper bound on the available escape routes. An optimal combination design must therefore maximize joint frequency while minimizing the unreached fraction of genes in the T-network. As shown in Figure 3 (top), prostate adenocarcinoma illustrates a clear trade-off: a panel of high-frequency genes achieves high joint frequency (0.69) but leaves ∼20% of the network unreached, whereas the perfect discriminatory panel reaches the whole population (f_T_ = 1) but leaves ∼30% of unreached nodes. In lung adenocarcinoma (LUAD), the same trade-off is much less pronounced (Figure 3, bottom), allowing a panel of two high-frequency T-genes to achieve both high coverage and low escape probability.

When the intervention targets NT-genes, an additional criterion must be considered: high frequency in normal samples (f_N_). Because NT genes are also active in normal tissues, those with high f_N_ are most relevant for normal tissue identity and, when reactivated, can most effectively restore normal differentiation programs. Thus, for NT targets, the optimization becomes three-dimensional: maximize f_T_, minimize unreached T-fraction, and prioritize high f_N_.

The probability of tumor escape after intervention is not determined solely by the fraction of the T-network that remains unreached. Our simulations reveal three interacting factors:

1. Fraction of the T-network reached by the reverse cascade (the complement of which defines the possible escape routes).
2. Tumor stage, measured by the initial number of active T-genes (younger tumors with fewer active genes are more plastic).
3. Activation rate of new T-genes, which represents the tumor’s adaptive response to the intervention. This rate can be influenced by the tumor’s intrinsic plasticity and by external factors such as compensatory signaling pathways.

The recovery time for younger tumors critically depends on the activation rate of T-genes. This underscores that effective therapy must not only maximize network reach but also suppress the tumor’s new activations rate, which is probably related to cellular division rate or genetic instability pathways.

The perspective presented in our paper aligns with emerging views on combination therapy and resistance. As Kauffman and colleagues have argued [3], multi-target strategies are essential for escaping entrenched cancer attractors. The NT-gene concept, combined with frequency analysis and the ratio of unreached genes, provides a rational basis for selecting such targets.

Several limitations should be acknowledged. First, the model is deliberately semi-quantitative and phenomenological. Second, causal relations inferred from expression data represent statistical dependencies, not necessarily direct mechanistic interactions; experimental validation remains essential. Third, the analysis uses bulk tissue expression, which may obscure intratumoral heterogeneity. Single-cell resolution data [31] will enable refined marker classification and frequency estimates.

In summary, a simple classification of genes based on their expression patterns across normal and tumor samples — coupled with basic concepts from deregulation network theory — yields testable predictions about the outcomes of genetic interventions. NT genes, particularly those with high activation frequency in both states, emerge as optimal targets for achieving at least partial phenotype reversal. The frequency metric provides a quantitative guide for target selection, while the unreached fraction of the T-network, tumor stage, and activation rate jointly determine escape risk. We hope this framework contributes to the rational design of next-generation cancer therapies aimed not merely at killing tumor cells but at restoring normal cellular identity.

## Acknowledgments

The authors acknowledge support by the Financial and International Projects Office of the Ministry of Sciences, Cuba (project PN692LH007-095).

## Author Contributions

GG created many of the ideas on which the methodology presented in the paper is based. RP stressed the possible role of NT-genes in therapy. AG designed the research plan and together with

RP wrote the initial draft. All authors analyzed and interpreted the results, contributed to the manuscript, and approved the final version.

